# Allele-specific transcript abundance: A pilot study in healthy centenarians

**DOI:** 10.1101/533398

**Authors:** Lauren C. Tindale, Nina Thiessen, Stephen Leach, Angela R. Brooks-Wilson

**Author notes:** Address correspondence to Angela Brooks-Wilson, PhD, Canada’s Michael Smith Genome Sciences Centre, BC Cancer Research Centre, 675 West 10th Ave, Vancouver, Canada, V5Z1L3. Current affiliation: Berlin Institute of Health, Charité Universitätsmedizin Berlin, Germany (Nina Thiessen).

## Abstract

The genetic basis of healthy aging and longevity remains largely unexplained. One hypothesis as to why long-lived individuals do not appear to have a lower number of common-complex disease variants, is that despite carrying risk variants, they express disease-linked alleles at a lower level than the wild-type alleles. Allele-specific abundance (ASA) is the different transcript abundance of the two haplotypes of a diploid individual. We sequenced the transcriptomes of four healthy centenarians and four mid-life controls. CIBERSORT was used to estimate blood cell fractions: neutrophils were the most abundant source of RNA, followed by CD8+ T cells, resting NK cells, and monocytes. ASA variants were more common in non-coding than coding regions. Centenarians and controls had a comparable distribution of ASA variants by predicted effect, and we did not observe an overall bias in expression towards major or minor alleles. Immune pathways were most highly represented among the gene set that showed ASA. Although we found evidence of ASA in disease-associated genes and transcription factors, we did not observe any differences in the pattern of expression between centenarians and controls in this small pilot study.

## Introduction

Several studies have shown that long-lived individuals do not appear to carry a substantially lower number of common-complex disease risk alleles compared to typical individuals (1-3). It may be that despite carrying risk alleles, they express the disease-related variants at a lower level. Allele-specific abundance (ASA) is the different transcript abundance of the two haplotypes of a diploid individual. ASA occurs across the genome in different biological processes and across multiple tissues (4). ASA can be caused by allele-specific alternative splicing, variation in transcriptional start or stop sites, *cis-*acting regulatory variants, epigenetic differences such as imprinting, DNA methylation, chromatin state, and differences in mRNA stability (4). ASA has been shown to be under genetic control in a study using lymphoblastoid cell lines (LCL), where monozygotic twins were found to have a more similar degree of ASA than unrelated controls (5).

Although different tissues have been shown to have important differences in expression, whole blood (WB) analysis has advantages. Peripheral blood is a cost-effective and clinically relevant tissue, it provides an overall view of the body since it comes into contact with almost all organs and tissues (6), and it is useful when looking for biomarkers for diseases where the tissue of interest is not readily available. Studies using WB collected in PAXgene tubes have identified and validated a gene expression signature that distinguished between Alzheimer’s Disease (AD) patients and cognitively healthy controls, showing that AD could be detected away from the primary site of the disease (7). Mutations causing expression differences in cardiac-restricted genes for Long QT Syndrome, Marfan Syndrome, and hypertrophic cardiomyopathy were also identified from WB from PAXgene tubes (8). WB is an appropriate tissue in which to assess overall patterns of ASE that may contribute to disease.

In a pilot study of four healthy centenarians and four mid-life controls, we hypothesized that the healthy centenarians may exhibit an overall pattern of lower expression of disease-associated alleles. This would effectively allow them to favour expression of non-risk alleles, despite carrying disease-causing variants. Heterozygous SNPs in the transcribed region of a gene may be used as indicators to determine if one copy of the gene is more highly expressed than the other. We tested whether preferential transcript abundance was observed, and whether it was skewed in centenarians versus controls by examining ASA in disease-associated genes and variants.

## Methods

### The Super-Seniors Study

Super-Seniors are aged 85 years and older with no reported history of cancer, cardiovascular disease (CVD), diabetes, major pulmonary disease, or dementia (9). Four Super-Senior centenarians aged 100-104 years, and four controls aged 50-56 years who were not selected for health, participated in a transcriptome pilot study. Research ethics board (REB) approval was received from the joint Clinical REB of BC Cancer and the University of British Columbia, and the REB of Simon Fraser University. Participants gave written informed consent.

### Sample preparation

WB was collected in PAXgene tubes, which preserve *in vivo* transcription levels. RNA was extracted using the PAXgene blood miRNA kit (QIAGEN) and globin depleted using the GLOBINclear™ kit (Thermo Fisher Scientific, Massachusetts, USA). Transcriptome and exome libraries were constructed using the Illumina TruSeq stranded mRNA kit and Agilent Technologies SureSelect v4+UTR kit, respectively.

### Analysis

CIBERSORT (10), an *in silico* flow cytometry tool, was used to estimate immune cell type abundances from gene expression data. The signature gene file of 22 distinct immune cell types provided by CIBERSORT was used as a reference for gene expression signatures.

Exome sequence calling was used to determine heterozygous SNPs, which were then matched to the transcriptome sequences. Autosomal variants were filtered for a read depth (RD) ≥30 and an alternate allele frequency >0.7 or <0.3.

Variants with ASA were compared by SNPEff predicted effect (11) and by major or minor allele bias as determined from minor allele frequencies reported in dbSNP (12). Gene lists were entered into QIAGEN Ingenuity^®^ Pathway Analysis (IPA) to determine enriched pathways and physiological functions. To look for disease associations we compared genes showing ASA to lists of publically available disease-associations. We looked for evidence of ASA in the 60 genes recommended by ACMG for reporting of incidental findings (13), as well as variants with SNP-trait associations that reached genome-wide significance listed in the NHGRI-EBI GWAS catalog v1.0.1 (14). Variants with ASA were input into SNP2TFBS to map SNPs to transcription factors (TF) (15).

## Results

CIBERSORT estimated the mean major cell types to be neutrophils, followed by CD8+ T cells, monocytes, and resting natural killer cells (Supplementary Figure S1 and Table S1).

After transcriptome alignment there were ~240-320 million mapped reads per sample. Centenarians had a mean of 344,167 (SD=14,369) variants per transcriptome and 130,068 (SD=10,188) heterozygous exome variants; controls had a mean of 355,220 (SD=136,358) variants per transcriptome and 128,775 (SD=3,029) heterozygous exome variants. There was a mean of 30,335 (SD=1,480) matches between the transcriptome and heterozygous exome variants in centenarians and 28,670 (SD=2,822) matches in controls. After filtering for RD ≥30, there were 16,679 (SD=1,108) matches in centenarians and 15,000 (SD=1,967) matches in controls. After filtering for alternate allele frequency >0.7 or <0.3, the final list of ASA variants contained 1,145 (SD=92) ASA variants in centenarians and 1,048 (SD=136) ASA variants in controls.

### Distribution and function of variants with ASA

Variants with ASA were most frequently non-coding: 3’UTR variants were most frequent, followed by intronic variants, downstream gene variants, and then coding missense and synonymous variants (Supplementary Figure S2).

There was no distinguishable difference in the proportion of variants with major allele bias versus minor allele bias when comparing centenarians and controls (Figure 1). Among ASA variants with an allelic ratio ≥0.7:0.3, centenarians had 49.1% of variants skewed to the minor allele, compared to 59.9% of variants in controls. At the more extreme ASA variants with an allelic ratio ≥0.9:0.1, centenarians had 53.3% of variants skewed to the minor allele, compared to 51.2% among controls. There was no difference in the proportion of variants skewed to the minor allele between centenarians and controls for allelic ratio ≥0.7:0.3 (X^2^=3.4; p=0.065) or for allelic ratio ≥0.9:0.1 (X^2^=3.33; p=0.068). There was no difference in the proportion in % skewed to the minor allele between the allelic ratio ≥0.7:0.3 and the allelic ratio ≥0.9:0.1 sets (X^2^=0.06; p=0.806).

**Figure 1.**
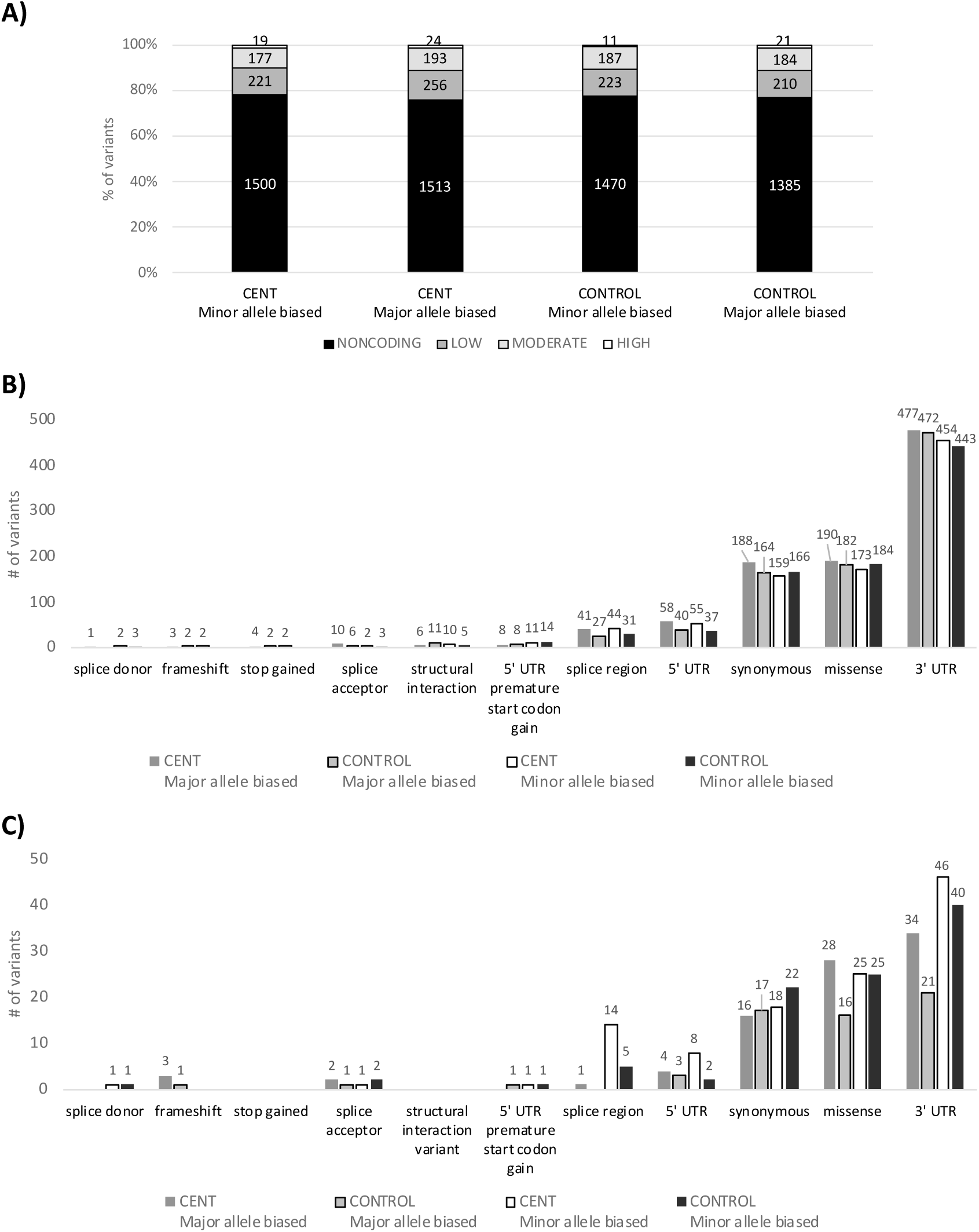
A) Proportion of major and minor allele bias by group. B) Coding region variants by major and minor allele bias (n=3700 coding variants with ASA ≥ 0.7:0.3). C) Coding region variants by major and minor allele bias with alternate allele frequency > 0.9 or < 0.1 (n=360 coding variants with ASA ≥ 0.9:0.1).

### Disease association of genes and variants with ASA

82 SNPs observed to have ASA ≥0.7:0.3 in either Super-Seniors or controls overlapped with SNPs in the NHGRI-EBI GWAS catalog. Trait associations from the catalog were filtered to keep only associations with diseases that were exclusion criteria for Super-Seniors. 21 disease-associated SNPs remained. Of these, 9 were skewed to the minor allele and 12 were skewed to the major allele. 7 were skewed towards the allele associated with disease, 9 were skewed away from the allele associated with disease, and 5 did not have a clear disease-associated allele (Table 1). 7 SNPs had ASA in multiple subjects, all of which were skewed in the same direction.

**Table 1.**
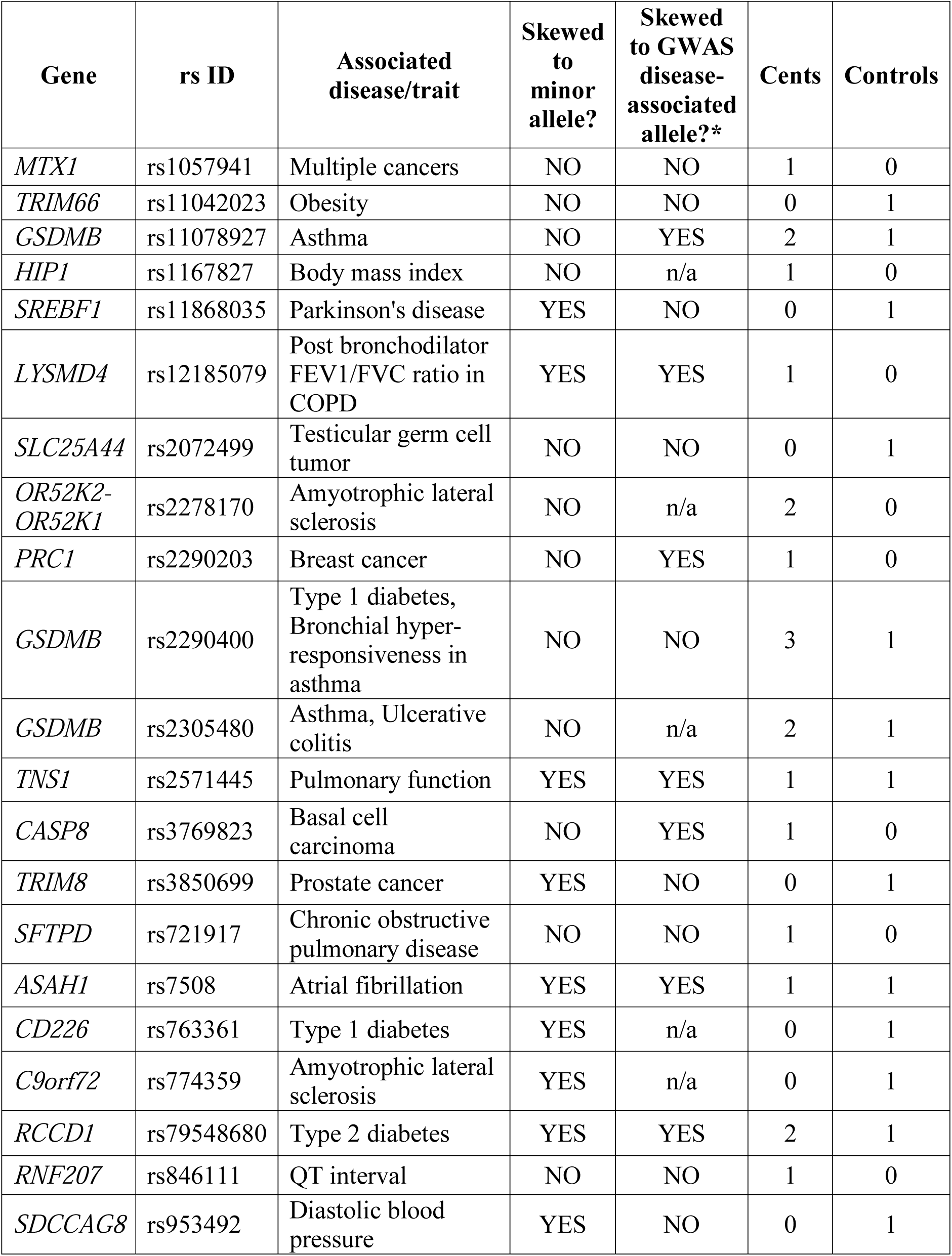
NHGRI-EBI GWAS catalog SNPs with evidence of allele-specific expression ≥ 0.7:0.3 in centenarians or controls.

There were 3373 genes where ASA was observed in either centenarians or controls. When the gene list was entered into QIAGEN IPA^®^ to determine enriched pathways and physiological functions, the top canonical pathways were: antigen presentation; natural killer cell signaling; CD28 signaling in T helper cells; type 1 diabetes mellitus signaling; and Th1 and Th2 activation.

Top centenarian genes with ASA were defined as genes for which there was evidence of ASA in 4 centenarians and only 1 or no controls, or in 3 centenarians and no controls; and vice versa for top genes with ASA in controls. There were 35 top genes with ASA in centenarians and 23 top genes with ASA in controls (Table S2). In QIAGEN IPA^®^ the top canonical pathways for the top genes with ASA in centenarians were, in order: citrulline-nitric oxide cycle; LXR/RXR activation; IL-12 signaling and production in macrophages; superpathway of citrulline metabolism; production of nitric oxide; and ROS in macrophages. The top canonical pathways for the top genes with ASA in controls were: LXR/RXR activation; role of osteoblast, osteoclasts and chondrocytes in rheumatoid arthritis; IL-10 signaling; CCR5 signalling in macrophages; and Fc-gamma Receptor-mediated phagocytosis in macrophages and monocytes.

Among the 60 ‘ACMG’ genes, we found that centenarians and controls showed ASA ≥0.7:0.3 in 8 and 11 genes, respectively (Table 2). Of note, in *LDLR* ASA was observed in 1 centenarian and all 4 controls; and in *TMEM43* ASA was observed in 3 centenarians and no controls. All other ASA variants in ACMG genes were unique with the exception of two SNPs in *PRKAG2*, which were each present in two subjects. One variant had a high predicted functional impact: a TTCT>TTCTCT indel at chr18:31068033 in *DSC2*.

**Table 2.** ACMG genes with allele-specific abundance ≥ 0.7:0.3 in centenarians or controls

| **Gene** | **Disease** | **Centenarians** | **Controls** |
| --- | --- | --- | --- |
| *APC* | Adenomatous polyposis coli |  | 1 |
| *BRCA1* | Breast-ovarian cancer, familial 1 |  | 1 |
| *DSC2* | Arrhythmogenic right ventricular cardiomyopathy, type 11 | 1 |  |
| *KCNH2* | Long QT syndrome 2 | 1 |  |
| *LDLR* | Familial hypercholesterolemia | 1 | 4 |
| *MYH11* | Aortic aneurysm, familial thoracic 4 | 1 |  |
| *NF2* | Neurofibromatosis, type 2 |  | 1 |
| *PRKAG2* | Familial hypertrophic cardiomyopathy 6 | 2 | 1 |
| *SDHAF2* | Paragangliomas 2 |  | 1 |
| *SMAD3* | Loeys-Dietz syndrome type 3 |  | 1 |
| *SMAD4* | Juvenile polyposis syndrome, | 1 |  |
| *TGFBR1* | Loeys-Dietz syndrome type 1A |  | 1 |
|  | Loeys-Dietz syndrome type 2A |  |  |
|  | Marfan's syndrome |  |  |
| *TMEM43* | Arrhythmogenic right ventricular cardiomyopathy, type 5 | 3 |  |
| *TNNI3* | Familial hypertrophic cardiomyopathy 7 |  | 1 |
| *TPM1* | Familial hypertrophic cardiomyopathy 3 |  | 1 |
| *TSC1* | Tuberous sclerosis 1 |  | 1 |
| *VHL* | Von Hippel-Lindau syndrome | 1 |  |
| **TOTAL** |  | **11** | **14** |

6039 unique variants with rsIDs were entered into SNP2TFBS (15). TF enrichment was determined by the ratio (total TF sites identified by SNP2TFBS): (TFs identified in the input list) (Supplementary Table S3). ZBTB33, Tcfcp2l1, NHLH1, TFAP2C, Pax2 were the top 5 most enriched TFs.

## Discussion

Greater RD increases the power to detect subtler ASA. Li et al. (16) calculated that at a RD of 30 there is ~60% statistical power to detect an allelic ratio of 0.7:0.3. An RD of 30 provides ~90% power to detect an allelic ratio of 0.8:0.2, and nearly 100% power to detect an allelic ratio of 0.9:0.1. They further calculated that to achieve RD of 30 in ~90% of exonic variants would require ~190 million reads, whereas to achieve RD of 30 in ~95% of exonic variants would require ~240 million reads. Since this is an exploratory study, to detect allelic ratios with a difference ≥0.7:0.3, samples were sequenced to 330-400 million raw reads per sample.

The main components of WB are red blood cells (RBC), platelets, and white blood cells. RBCs contribute globin transcripts which have been found to make up on average 60% of the mRNA transcripts of WB; however, after globin reduction this decreases to 0.1-0.4% (6). White blood cells are comprised of numerous cell types including neutrophils, eosinophils, basophils, lymphocytes, and monocytes. Some studies utilize LCLs immortalized by Epstein-Barr virus (EBV), but gene expression has been shown to vary greatly between LCL and WB samples (17). While LCLs are simpler to interpret because they represent a single cell type, they may be less likely to reflect *in vivo* expression levels due to expression changes during EBV transformation (18) and degradation between cell collection and RNA preparation (19).

Overall, we found that centenarians and controls had a comparable distribution of ASA variants with no difference in proportion by predicted effect. ASA variants were more common in non-coding than coding regions. We did not observe an overall bias in expression towards the major or minor alleles in centenarians or controls.

Immune pathways were most highly represented among the gene set that showed ASA. Immunosenescence is seen in the adaptive as well as innate immune systems (20). The occurrence of immune dysfunction with aging is well established, including increased susceptibility to infection, frequency of neoplasia, inflammation, and autoimmune response (21). A common factor underlying the pathogenesis of many age-related chronic diseases including CVD, cancer, neurodegenerative diseases, and diabetes is inflammation (20). It has been previously suggested that centenarians may have genetic factors that allow them to better maintain immune function as they age (22,23).

When looking at ACMG genes with potential clinical utility (13), we found evidence of ASA in some genes related to disease etiology. Although it may be an artifact of small sample size, *LDLR*, for which ASA was observed in 1 centenarian and all 4 controls, and *TMEM43*, for which ASA was observed in 3 centenarians and no controls, are potentially interesting to look at in a larger study. *LDLR* is associated with familial hypercholesterolemia and is a major apoE receptor; variants of the *APOE* gene are the most replicated association with longevity and healthy aging phenotypes (24) including the Super-Senior phenotype (25). *TMEM43* is associated with arrhythmogenic right ventricular cardiomyopathy, type 5. ASA variants also overlapped with disease-associated NHGRI-EBI GWAS catalogue variants. It is possible that among patients with disorders associated with these genes, even when a pathogenic variant is not identified, ASA may be playing a role in disease pathogenicity if a *cis*-acting factor is affecting gene expression. There was evidence that ASA variants overlapped with numerous TFs, which are possible mechanisms for *cis*-acting affects.

A caveat to the ASA that we observed is that it may not reflect true *in vivo* RNA expression levels. As well, ASA can be caused by numerous mechanisms that cannot be distinguished in the present study. It is possible that some of our observed cases of ASA could be the result of alternative transcripts that have been collapsed into single loci.

As a pilot study, the sample size was very small, and may lead to false negative findings. We did not find evidence for a pattern of lower expression of disease-associated variants in centenarians compared to mid-life controls. The presence of ASA in some ACMG genes suggests that in patients with disorders that are highly associated with a certain gene, but for which no causal variant is apparent, looking at ASA may be of interest. As well, the representation of immune pathways among the ASA gene set of centenarians may be related to decreased immune function with aging.

## Acknowledgements

Study design by LCT, ABW; lab work by LCT, SL; analysis by LCT, NT; draft of manuscript by LCT; all authors read and approved the manuscript

