## Supplementary files for "Allele-specific transcript abundance: A pilot study in healthy centenarians"

**Supplementary Data**

**Index**


Supplementary Table S1. Estimated average immune cell type fractions across all 8 samples

Supplementary Table S2. List of 35 top genes that show ASA in centenarians and 23 top genes that show ASA in controls. These are genes for which there was evidence of ASA in 4 centenarians and 1 or no controls, or in 3 centenarians and no controls; or vice versa for controls

Supplementary Table S3. Results of mapping SNPs with allele-specific abundance to transcription factors using SNP2TFBS. Table of top 50 transcription factor enrichment statistics produced by SNP2TFBS (ccg.vital-it.ch/snp2tfbs/).

Supplementary Figure S1. Proportion of immune cell types as estimated by CIBERSORT, by sample. Figure produced using CIBERSORT (cibersort.stanford.edu).

Supplementary Figure S2. Proportion of variants showing allele-specific expression, by predicted effect.

Supplementary Table S1. Estimated average immune cell type fractions across all 8 samples

| Cell type | Average relative % | Average relative call fraction | SD | Min | Max |
|---|---|---|---|---|---|
| Neutrophils | 42.37% | 0.42 | 0.19 | 0.19 | 0.74 |
| T cells CD8 | 17.21% | 0.17 | 0.13 | 0.022 | 0.44 |
| NK cells resting | 12.30% | 0.12 | 0.072 | 0.050 | 0.23 |
| Monocytes | 9.08% | 0.091 | 0.068 | 0.0026 | 0.20 |
| T cells regulatory (Tregs) | 4.36% | 0.044 | 0.023 | 0.013 | 0.081 |
| B cells naive | 3.54% | 0.035 | 0.036 | 0 | 0.092 |
| Mast cells resting | 3.44% | 0.034 | 0.011 | 0.013 | 0.048 |
| T cells CD4 memory resting | 2.51% | 0.025 | 0.025 | 0 | 0.077 |
| T cells CD4 memory activated | 1.88% | 0.019 | 0.015 | 0.0012 | 0.046 |
| B cells memory | 1.25% | 0.012 | 0.015 | 0 | 0.034 |
| T cells CD4 naive | 0.63% | 0.0063 | 0.012 | 0 | 0.036 |
| Macrophages M0 | 0.46% | 0.0046 | 0.0079 | 0 | 0.022 |
| NK cells activated | 0.43% | 0.0043 | 0.0069 | 0 | 0.016 |
| Plasma cells | 0.38% | 0.0038 | 0.0036 | 0 | 0.0097 |
| Dendritic cells resting | 0.11% | 0.0011 | 0.0016 | 0 | 0.0037 |
| T cells follicular helper | 0.05% | 0.00048 | 0.0014 | 0 | 0.0039 |
| Dendritic cells activated | 0.01% | 0.000050 | 0.00014 | 0 | 0.00040 |
| T cells gamma delta | 0.00% | 0 | 0 | 0 | 0 |
| Macrophages M1 | 0.00% | 0 | 0 | 0 | 0 |
| Macrophages M2 | 0.00% | 0 | 0 | 0 | 0 |
| Mast cells activated | 0.00% | 0 | 0 | 0 | 0 |
| Eosinophils | 0.00% | 0 | 0 | 0 | 0 |

Immune cell types were determined using CIBERSORT. The signature gene file of 22 immune cell types provided by CIBERSORT was used as a reference for gene expression signatures. Table produced using CIBERSORT (cibersort.stanford.edu).

Supplementary Table S2. List of 35 top genes that show ASA in centenarians and 23 top genes that show ASA in controls. These are genes for which there was evidence of ASA in 4 centenarians and 1 or no controls, or in 3 centenarians and no controls; or vice versa for controls

| **Subjects with ASE** | **Gene** |
|---|---|
| 4 Cents - 0 Controls | LINC01060 |
| 4 Cents - 0 Controls | LINC01262 |
| 4 Cents - 0 Controls | NOS2 |
| 4 Cents - 0 Controls | TCF25 |
| 3 Cents - 0 Controls | ATF7IP |
| 3 Cents - 0 Controls | C3orf58 |
| 3 Cents - 0 Controls | CAMK2N1 |
| 3 Cents - 0 Controls | CD151 |
| 3 Cents - 0 Controls | CEP295 |
| 3 Cents - 0 Controls | EIF4G2 |
| 3 Cents - 0 Controls | ELMSAN1 |
| 3 Cents - 0 Controls | FES |
| 3 Cents - 0 Controls | JAML |
| 3 Cents - 0 Controls | LILRA1 |
| 3 Cents - 0 Controls | LINC00226 |
| 3 Cents - 0 Controls | LINC00221 |
| 3 Cents - 0 Controls | LLGL2 |
| 3 Cents - 0 Controls | LPIN1 |
| 3 Cents - 0 Controls | MED16 |
| 3 Cents - 0 Controls | MEGF6 |
| 3 Cents - 0 Controls | NELFCD |
| 3 Cents - 0 Controls | ORM1 |
| 3 Cents - 0 Controls | PDCD6IP |
| 3 Cents - 0 Controls | RNF44 |
| 3 Cents - 0 Controls | RRN3P2 |
| 3 Cents - 0 Controls | SCRN1 |
| 3 Cents - 0 Controls | TMEM43 |
| 3 Cents - 0 Controls | TRIM39 |
| 3 Cents - 0 Controls | UNC13D |
| 3 Cents - 0 Controls | WBP2 |

| **Subjects with ASE** | **Gene** |
|---|---|
| 3 Cents - 0 Controls | ZFP57 |
| 3 Cents - 0 Controls | ZNF718 |
| 4 Cents - 1 Control | KRT72 |
| 4 Cents - 1 Control | SNORA10 |
| 4 Cents - 1 Control | TPTE2P5 |
| 4 Cents - 1 Control | TRG-AS1 |
| 0 Cents - 4 Controls | WDR90 |
| 1 Cent - 4 Controls | DLGAP4 |
| 1 Cent - 4 Controls | IL18RAP |
| 1 Cent - 4 Controls | LDLR |
| 1 Cent - 4 Controls | LDOC1L |
| 0 Cents - 3 Controls | ANXA5 |
| 0 Cents - 3 Controls | COX5BP7 |
| 0 Cents - 3 Controls | CARD8 |
| 0 Cents - 3 Controls | CD247 |
| 0 Cents - 3 Controls | CDC42EP1 |
| 0 Cents - 3 Controls | CPA5 |
| 0 Cents - 3 Controls | FAM118A |
| 0 Cents - 3 Controls | PSPHP1 |
| 0 Cents - 3 Controls | FYB |
| 0 Cents - 3 Controls | HSD17B1 |
| 0 Cents - 3 Controls | HVCN1 |
| 0 Cents - 3 Controls | IL1RN |
| 0 Cents - 3 Controls | MFSD9 |
| 0 Cents - 3 Controls | NOC4L |
| 0 Cents - 3 Controls | PTK2B |
| 0 Cents - 3 Controls | SGSH |
| 0 Cents - 3 Controls | SNX22 |
| 0 Cents - 3 Controls | UPK3A |
| 0 Cents - 3 Controls | VNN1 |

Supplementary Table S3. Results of mapping SNPs with allele-specific abundance to transcription factors using SNP2TFBS. Table of top 50 transcription factor enrichment statistics produced by SNP2TFBS (ccg.vital-it.ch/snp2tfbs/).

| **TF Name** | **TF-SNP matches (genome-wide)** | **TF-SNP hits (from query)** | **Fraction of TF-SNP hits** | **Enrichment** | **P-value** |
|---|---|---|---|---|---|
| ZBTB33 | 2855 | 5 | 0.00175 | 4.49 | 0.0057 |
| Tcfcp2l1 | 10304 | 16 | 0.00155 | 3.98 | 4.980E-06 |
| NHLH1 | 12332 | 17 | 0.00138 | 3.53 | 1.180E-05 |
| TFAP2C | 28651 | 38 | 0.00133 | 3.40 | 2.000E-10 |
| Pax2 | 6140 | 8 | 0.00130 | 3.34 | 0.0033 |
| Mafb | 6619 | 8 | 0.00121 | 3.10 | 0.0051 |
| Tcf12 | 11250 | 13 | 0.00116 | 2.96 | 6.301E-04 |
| KLF5 | 61743 | 70 | 0.00113 | 2.90 | 1.070E-14 |
| TFAP2A | 23552 | 26 | 0.00110 | 2.83 | 3.840E-06 |
| Myb | 5513 | 6 | 0.00109 | 2.79 | 0.0226 |
| REST | 19562 | 21 | 0.00107 | 2.75 | 4.660E-05 |
| EGR1 | 70916 | 75 | 0.00106 | 2.71 | 3.730E-14 |
| Myog | 13245 | 14 | 0.00106 | 2.71 | 9.299E-04 |
| E2F1 | 16499 | 17 | 0.00103 | 2.64 | 3.735E-04 |
| SP2 | 111891 | 115 | 0.00103 | 2.63 | 3.430E-20 |
| THAP1 | 11687 | 12 | 0.00103 | 2.63 | 0.0026 |
| E2F3 | 39019 | 40 | 0.00103 | 2.63 | 7.620E-08 |
| Myod1 | 16791 | 17 | 0.00101 | 2.59 | 4.535E-04 |
| SP1 | 98565 | 93 | 0.00094 | 2.42 | 2.370E-14 |
| CTCF | 19639 | 18 | 0.00092 | 2.35 | 9.679E-04 |
| Tcf3 | 23038 | 21 | 0.00091 | 2.34 | 4.127E-04 |
| Erg | 14483 | 13 | 0.00090 | 2.30 | 0.0054 |
| Nkx3-2 | 4504 | 4 | 0.00089 | 2.28 | 0.1020 |
| NRF1 | 10231 | 9 | 0.00088 | 2.25 | 0.0210 |
| HNF4A | 11861 | 10 | 0.00084 | 2.16 | 0.0201 |
| PPARG_RXRA | 21593 | 18 | 0.00083 | 2.14 | 0.0027 |
| FLI1 | 13204 | 11 | 0.00083 | 2.13 | 0.0165 |
| ESR1 | 24743 | 20 | 0.00081 | 2.07 | 0.0023 |
| E2F6 | 33466 | 27 | 0.00081 | 2.07 | 4.519E-04 |
| PLAG1 | 34341 | 27 | 0.00079 | 2.01 | 6.605E-04 |
| NR2C2 | 19125 | 15 | 0.00078 | 2.01 | 0.0097 |
| MAFF | 10436 | 8 | 0.00077 | 1.96 | 0.0554 |
| MAFK | 9281 | 7 | 0.00075 | 1.93 | 0.0749 |
| Meis1 | 13726 | 10 | 0.00073 | 1.87 | 0.0464 |

| | | | | | |
|---|---|---|---|---|---|
| Ets1 | 22116 | 16 | 0.00072 | 1.85 | 0.0154 |
| HNF4G | 11382 | 8 | 0.00070 | 1.80 | 0.0816 |
| T | 4282 | 3 | 0.00070 | 1.79 | 0.2352 |
| JUN | 7179 | 5 | 0.00070 | 1.78 | 0.1525 |
| E2F4 | 14380 | 10 | 0.00070 | 1.78 | 0.0595 |
| Hltf | 4415 | 3 | 0.00068 | 1.74 | 0.2489 |
| ESR2 | 12522 | 8 | 0.00064 | 1.64 | 0.1216 |
| JUNB | 6262 | 4 | 0.00064 | 1.64 | 0.2305 |
| MZF1_1-4 | 126853 | 81 | 0.00064 | 1.64 | 1.690E-05 |
| INSM1 | 17275 | 11 | 0.00064 | 1.63 | 0.0808 |
| Bach1_Mafk | 11016 | 7 | 0.00064 | 1.63 | 0.1439 |
| Klf4 | 50628 | 32 | 0.00063 | 1.62 | 0.0065 |
| Arnt_Ahr | 75959 | 48 | 0.00063 | 1.62 | 0.0010 |
| Rxra | 9788 | 6 | 0.00061 | 1.57 | 0.1873 |
| RXR_RAR_DR5 | 10022 | 6 | 0.00060 | 1.53 | 0.2010 |
| Hand1_Tcfe2a | 6794 | 4 | 0.00059 | 1.51 | 0.2753 |

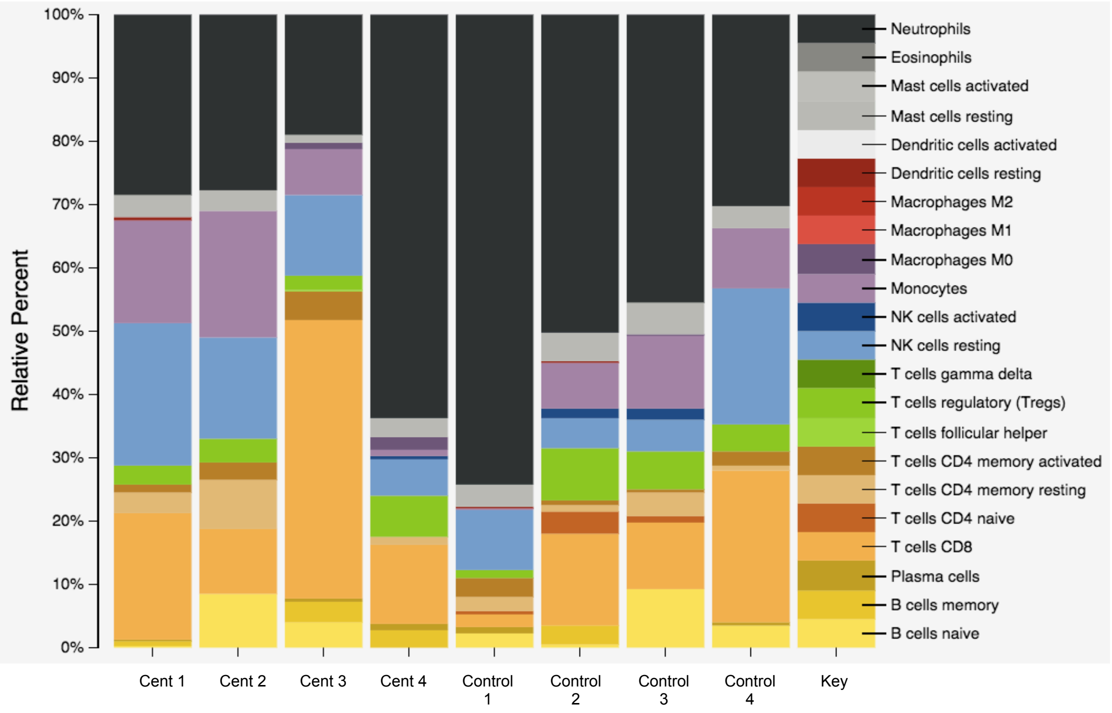


Supplementary Figure S1. Proportion of immune cell types as estimated by CIBERSORT, by sample. Figure produced using CIBERSORT (cibersort.stanford.edu).

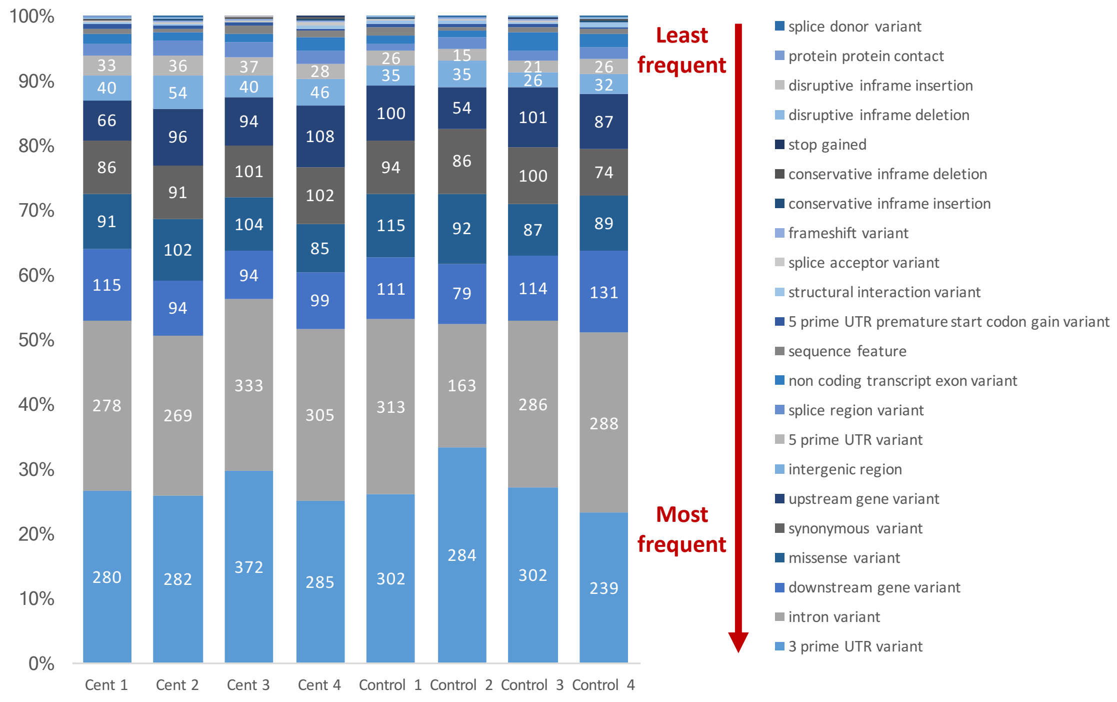


Supplementary Figure S2. Proportion of variants showing allele-specific expression, by predicted effect.
